# Proxy-based model to assess the relative contribution of ballast water and biofouling’s potential propagule pressure and prioritize vessel inspections

**DOI:** 10.1101/2021.02.10.430588

**Authors:** Lina Ceballos-Osuna, Chris Scianni, Maurya Falkner, Raya Nedelcheva, Whitman Miller

**Affiliations:** Marine Invasive Species Program, California State Lands Commission, Sacramento, CA, 95825, USA; Smithsonian Environmental Research Center, Edgewater MD, 21037, USA

## Abstract

Commercial shipping is the primary pathway of introduction for aquatic nonindigenous species, mainly through the mechanisms of ballast water and biofouling. In response to this threat, regulatory programs have been established across the globe to regulate and monitor commercial merchant and passenger vessels to assess compliance with local requirements to reduce the likelihood of NIS introductions. Resource limitations often determine the inspection efforts applied by these regulatory agencies to reduce NIS introductions. We present a simple and adaptable model that prioritizes vessel arrivals for inspection using proxies for potential propagule pressure, namely a ships’ wetted surface area as a proxy for the likelihood of biofouling-mediated potential propagule pressure and ballast water discharge volume as a proxy for ballast water-mediated potential propagule pressure. We used a California-specific dataset of vessels that arrived at California ports between 2015 and 2018 to test the proposed model and demonstrate how a finite set of inspection resources can be applied to target vessels with the greatest potential propagule pressure. The proposed tool is adaptable by jurisdiction, scalable to different segments of the vessel population, adjustable based on the vector of interest, and versatile because it allows combined or separate analyses of the PPP components. The approach can be adopted in any jurisdiction across the globe, especially jurisdictions without access to, or authority to collect, risk profiling data or direct measurements for all incoming vessel arrivals.

## Introduction

Commercial shipping is the primary pathway of introduction for aquatic nonindigenous species (NIS) that have established within coastal and estuarine waters globally [1-4]. Commercial vessels transport NIS from one location to another through the uptake and discharge of ballast water and as a result of biofouling associated with the submerged portions of the hull, or the vessel’s wetted surface area (WSA) [5-7].

Ballast water is taken on board a vessel for trim and stability purposes and to offset the mass imbalance of the vessel during cargo loading and unloading operations. However, when ballast water is taken on board in one location, planktonic communities are inadvertently entrained, along with sediments and benthic biota that may have been suspended in the water column by vessel activity. NIS are released when the ballast water is eventually discharged in another port.

Biofouling refers to the biota attached to, or associated with, the submerged portions of a vessel (i.e., the WSA) [8]. Biofouling organisms can be sessile or mobile, and range in size from microscopic bacterial biofilms to larger macrofauna. These organisms can either cling to the vessel’s wetted surfaces (e.g., biofilms, barnacles, bivalves, bryozoans, algae) or find refuge in internal cavities where they are protected from shear forces at the hull-water interface (e.g., mobile crustaceans, fish). The biofouling communities travel wherever the vessel travels, transporting organisms to new ports and places. These biofouling organisms can be released as adults, larvae, or other propagules, either through physical forces such as the hull rubbing on a pier piling or displacement from an activated bow thruster, or by natural release and spawning.

In response to the biosecurity threat posed by commercial shipping, jurisdictions across the globe have established (or are in the process of establishing) regulatory programs tasked with reducing the likelihood of NIS introductions via ballast water and biofouling (e.g., California’s Marine Invasive Species Program). Each of these local, state, federal, and international programs have the goal of implementing ballast water or biofouling management requirements/regulations to prevent species introductions in their respective jurisdictions. A critical element of these efforts is a robust outreach and inspection program to assess and improve compliance. Active communication between regulators (e.g., Port State Control) and a vessel’s crew is the most straightforward way of getting location-specific requirements into the hands of the people directly responsible for ballast water and biofouling management actions. Ideally, all arriving vessels would be boarded for outreach and inspection, but resource limitations such as staffing levels and funding often require regulatory agencies to make decisions about which vessels to board and which vessels to bypass. How agencies allocate their limited resources to inspections can vary along a spectrum from arbitrary to deliberate. Using a standardized, data-driven prioritization approach designed for conditions specific to each jurisdiction could greatly improve the decision-making process.

A variety of risk-based vessel prioritization approaches have been described in recent years [9-11], typically with the goal of assessing the likelihood of a vessel introducing NIS through ballast water or biofouling. Such approaches and their reliability differ depending on the quantity and quality of input data available (Fig 1). Agencies with detailed and readily accessible data on ballast water and/or biofouling operational and maintenance practices, or with detailed source and recipient port information, are in the strongest position to use intricate risk-based prioritization approaches for boarding and inspecting vessels to assess compliance. However, other approaches based on available proxies can also be useful when detailed vessel operational or environmental matching data are limited or not readily accessible.

**Fig 1.**
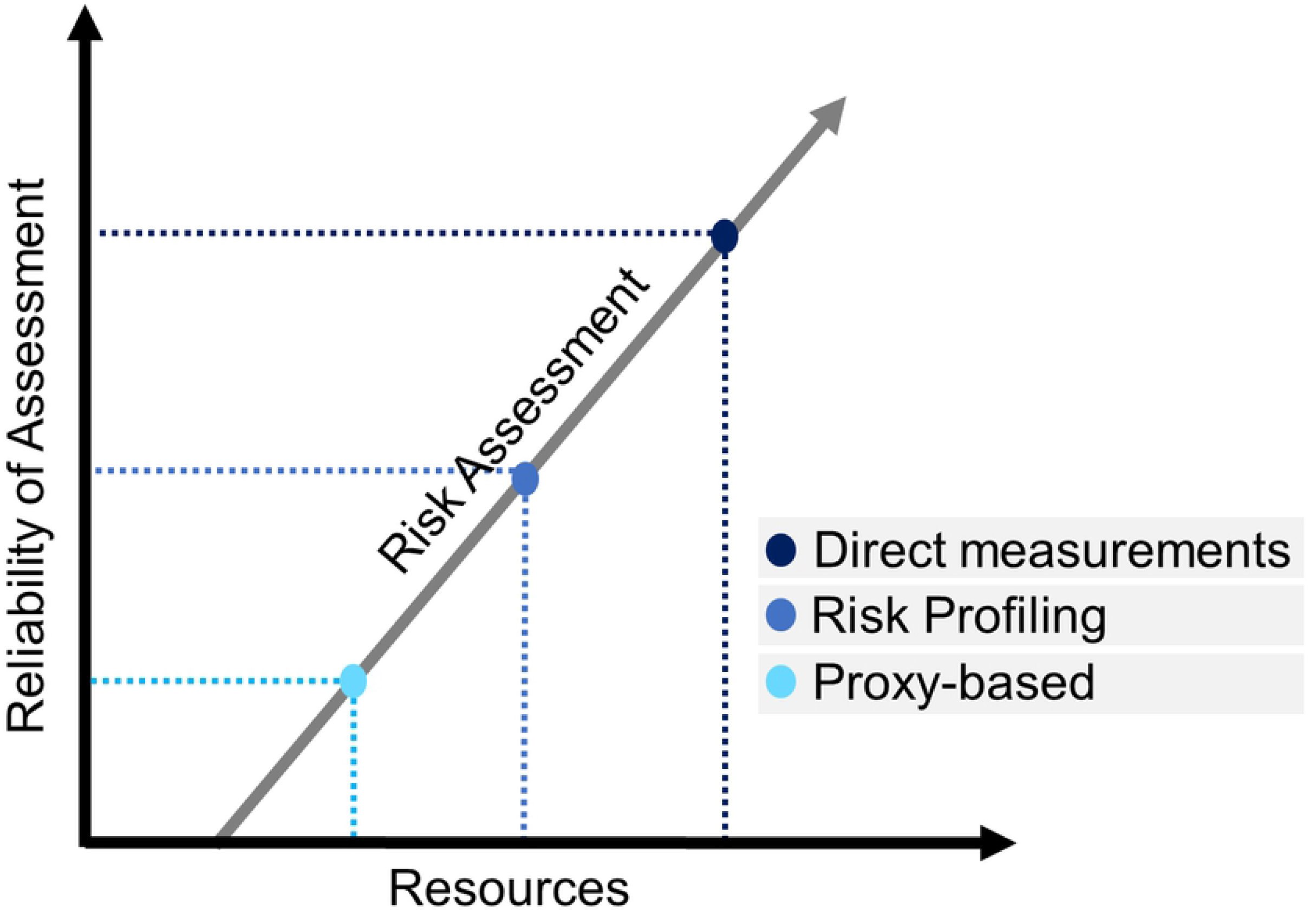
Conceptual approaches for risk-based prioritization. Reliability of an approach is dependent on the quantity and quality of available resources. Direct measurement approaches may rely on physical sampling of ballast water, surveying of a vessel’s wetted surface area, or similar methods. Risk profiling approaches may rely on environmental matching, vessel operational profile, or other similar methods. Proxy-based approaches may rely on ballast water discharge volumes and wetted surface area as potential propagule pressure parameters, or other appropriate measurements.

In this paper, we present a proxy-based model that prioritizes vessel arrivals for inspection using fundamental information about ships’ WSA and BWD volumes to enable agencies to make more protective data-driven decisions when resources are limiting.

Despite the low numbers of empirical studies able to prove the direct relationship between vector activity and invasion success [12], the positive relationship between propagule supply (e.g., the release of organisms in ballast water discharged or from the WSA of the vessel) and the likelihood of introductions is well recognized [13, 14]. Based on the assumptions that ballast water and biofouling are managed consistent with local requirements and that the likelihood of introduction increases with increasing propagule supply (also recognizing the uncertainty of translating proxy-based parameters into invasion success, [12, 15]), we present a reliable and simplified alternative model to prioritize vessel arrivals based on potential propagule supply (i.e., potential propagule pressure, PPP) for jurisdictions with limited resources.

The model is based on PPP proxies, as described in Lo et al. [16], that are readily available to most jurisdictions. The proxies used in this approach are BWD volume as a proxy for ballast water PPP and WSA as a proxy for biofouling PPP. The targeted users for this approach are jurisdictions that do not have access to detailed vessel operational or environmental matching data between source and recipient ports, or that do not have the capacity and time to make direct measurements.

An additional benefit of this model is that it allows for identification of the relative pressure of each component of the overall PPP, either combined or independently (e.g., by location, vessel type, and/or vector). From a management perspective, it could be particularly useful when assessing cumulative supply pressure across different ports to guide decisions about where limited resources should be allocated and target those ports that proportionally receive the greatest PPP.

To test the proposed model, we used a California-specific dataset from vessels that arrived at California ports between 2015 and 2018 (S1 Dataset) and compared with California’s current prioritization scheme that allocates available resources to meet the legislative mandate of inspecting 25% of the vessels arriving at California ports to assess compliance with California’s ballast water and biofouling management requirements.

## Methods

### Wetted surface area

The use of WSA as a proxy for PPP relies on the assumption that the likelihood of introduction increases as the area of colonizable surface, including niche areas (e.g., sea chests, rudders), increases. We used the same vessel dataset as Miller *et al*. [7], where the WSA for 373,833 vessel arrivals at United States ports was calculated using the WSA equation and the coefficients reported by Van Maanen and Van Oossanen [17]. A relationship between WSA and Gross Tonnage (GT) was established via regression analysis (Table 1) for each of the following vessel-types: General Cargo, Passenger, RO-RO (Auto carriers), Bulker, Container, and Tanker. GT was used here, rather than Net Tonnage as used by Miller *et al*. [7], because it is a more readily available metric and to match with an existing California-specific vessel dataset used to trial the model. Unmanned barges (including their respective tug) and articulated tug-barges (ATB) were not included in the regression analysis due to the variability within these groups, but their WSA was calculated directly for each vessel following Van Maanen and Van Oossanen [17] coefficients.

**Table 1.** Calculation of wetted surface area. Regression models for specific vessel types used to estimate the wetted surface area (WSA) from gross tonnage (GT). Niche proportion [18] represents the fraction of a vessel’s WSA that is accounted for by niche areas (e.g., sea chests, rudders).

| | Regression Equation*<br>$WSA = m(GT)^b$ | n** | r<sup>2</sup> | Niche Proportion (N<sub>p</sub>) |
| --- | --- | --- | --- | --- |
| General | WSA=20.02(GT)<sup>0.5728</sup> | 4172 | 0.907 | 0.09 |
| Passenger | WSA =5.46 (GT)<sup>0.6951</sup> | 2659 | 0.996 | 0.27 |
| RO-RO | WSA =25.04 (GT)<sup>0.5309</sup> | 1929 | 0.901 | 0.09 |
| Bulker | WSA =15 (GT)<sup>0.6294</sup> | 8804 | 0.988 | 0.07 |
| Container | WSA =10.66 (GT)<sup>0.6501</sup> | 3065 | 0.984 | 0.09 |
| Tanker | WSA =17.57 (GT)<sup>0.6105</sup> | 8292 | 0.988 | 0.08 |
| ATB | *Calculated per vessel* |  |  | 0.033 (barge) + 0.25<br>(tug) |
| Unmanned<br>barges + Tug | *Calculated per vessel* |  |  | 0.033 (barge) + 0.25<br>(tug) |

\*Regressions are based on first order of polynomial relationships. *m*= slope, *b*= intercept
\*\* Number of vessels used in the regression model

In addition to creating regression equations to estimate WSA using GT, we also applied the estimated proportion of a vessel’s WSA accounted for by niche areas, as reported by Moser *et al*. [18] for each vessel type (Table 1). Niche proportions for ATBs and unmanned barges and their tugs were also estimated using specific niche area values reported by Moser *et al*. [18] and adding all niche areas expected for typical ATBs, unmanned barges, and tugs (K. Reynolds, *pers comm*, 2020). These two metrics, estimated WSA and niche proportion (N_p_), can be used to calculate total WSA (TWSA) using equation 1:

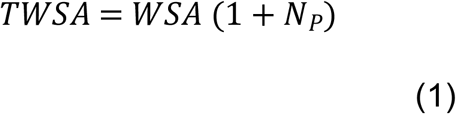

(see supplementary material for the step-by-step process (S2), the R script (S3), and the data frame template (S4) to calculate TWSA)

### Ballast water discharge volume

The use of BWD volume as a proxy for PPP relies on the assumptions that propagule supply increases as BWD volume increases [16] and that management is consistently applied in compliance with local requirements. We recognize the limitations that this assumption may have when trying to predict invasion success [12, 14, 15], however, our intent is not to measure probability of species establishment. Instead, our intention is to rely on the positive relationship between BWD volume and propagule supply [12, 13] to prioritize limited resources with the goal of assessing vessel compliance to reduce the likelihood of introduction using readily available information.

BWD volume data are available to most jurisdictions in the form of ballast management reporting for each vessel arrival. In the U.S., the National Ballast Information Clearinghouse (NBIC) provides vetted BWD and management information for all U.S. arrivals to State and Territorial agencies via an online public data portal (https://nbic.si.edu/). Other factors like water origin, environmental matching between source and recipient waters, and management strategy also influence NIS introductions. However, this information is not readily available in most cases, and can be complex and challenging to analyze. As resources increase, additional factors can be included to improve reliability of the analysis (Fig 1).

### Potential propagule pressure (PPP): Combined influence of ballast water discharge and biofouling

We describe a proxy-based model to calculate PPP scores using TWSA and BWD volume to prioritize vessel arrivals for inspection, targeting vessels that are more likely to introduce NIS at a specific location, assuming that all vessels have managed consistent with local requirements. This approach can also help identify the ports that receive more PPP due to the frequency of arrivals.

For jurisdictions that have minimal resources to accomplish this, individual vessel PPP scoring will allow a simple prioritization scheme specifically designed for their own population of vessels. The process requires a representative number of historical arrivals (e.g., one month, one year, multiple years, referred to as “Population data” in S4) to identify the maximum individual vessel value of both TWSA (i.e., _*max*_*TWSA*_*ind*_), calculated using the regression equations in Table 1 for each vessel type and Equation 1, and BWD (i.e., _*max*_*BWD*_*ind*_) from vessel reported data. A PPP score for each new vessel arrival can then be calculated using the estimated TWSA (i.e., *TWSA*_*ind*_) and the BWD volume (i.e., *BWD*_*ind*_) specific for that arrival as the input values in Equation 2.

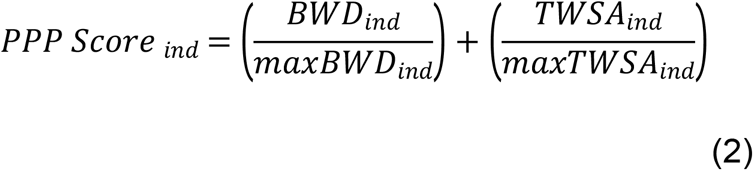

(see the R script provided in S3 to calculate PPP score)

Each vessel arrival will have an individual vessel PPP score that can be used to sort and prioritize arrivals relative to the other vessels in the population.

#### PPP scores – Component parts and cumulative scores

The relative contribution of each PPP component (i.e., ballast water and biofouling) can be calculated separately using either of the parenthetical components on the right-hand side of Equation 2. The overall influence of either ballast water or biofouling on PPP over time (i.e., incorporating frequency of arrival) or geographical region (i.e., incorporating all arrivals for specific ports) can then be added appropriately. Similarly, the total PPP score can be added cumulatively across a region or over certain time periods to compare between ports or over time. Likewise, cumulative scores can be assessed by vessel type or a myriad of other categories.

Because conducting management to meet local requirements is considered the first layer to reduce the likelihood of NIS introductions, the proposed approach assumes that each jurisdiction already has voluntary or mandatory management requirements for both ballast water (e.g., ballast water exchange or ballast water treatment systems to meet discharge performance standards) and biofouling (e.g., antifouling coatings) in place. All vessels evaluated under the PPP proxy-based prioritization scheme are assumed to be compliant with local management requirements; if they are not, they should be automatically categorized as high priority for inspection independent of the estimated PPP.

### Model trial using California data

We used data from four years of arrivals at California ports (2015-2018, S1 dataset) to trial the proposed model (For clarification, this proposed approach is not the current vessel inspection prioritization scheme used in California). BWD volume data were obtained from Ballast Water Management Reports submitted to the California State Lands Commission and vessel arrival data were obtained from the Marine Exchanges of Southern California and the San Francisco Bay Region.

#### PPP scoring by individual vessel - California

Using the TWSA and BWD volume data for each arriving vessel during the first three years of our dataset (2015-2017), we identified the maximum values for BWD (i.e., _*max*_*BWD*_*ind*_) and TWSA (i.e., _*max*_*TWSA*_*ind*_) for future use in Equation 2. We then used the TWSA and BWD volume data for each arriving vessel during the final year of our dataset (2018) in Equation 2 with the maximum values identified earlier. These calculations produced individual vessel PPP scores for each arrival during 2018 to evaluate how vessels would be prioritized daily according to each arriving vessel’s relative PPP.

Currently, California has a mandate to inspect at least 25% of all arrivals, corresponding to an average of 6 vessels inspected per day. Using this number as the target for high priority arrivals to inspect (and representing the resources available for the trial), we identified the 6 greatest PPP scores per day and analyzed how each vessel type would be represented under the arrivals prioritized for inspection.

#### Cumulative PPP scores (by location and vessel type)

To demonstrate how cumulative PPP scores can be used to inform the distribution of resources, we added all 2018 PPP scores by vessel type and by port.

## Results

### Model results using California data

#### PPP scores by individual vessel in California

Each component of the individual vessel PPP score (i.e., the two parenthetical values on the right-hand side of Equation 2) is expected to produce unitless values within the range between 0 and 1. Therefore, the combined PPP score_ind_ for each vessel is expected to range between 0 and 2. California’s 2018 dataset produced individual vessel PPP scores ranging from 0.1 to 1.5. Passenger (0.60 ± 0.005 SE) and Container (0.55 ± 0.003 SE) vessels accounted for the greatest mean PPP scores, whereas

General cargo vessels (0.22 ± 0.007 SE) and Unmanned Barges (0.11 ± 0.001 SE) exhibited the lowest mean values (Fig 2).

**Fig 2.**
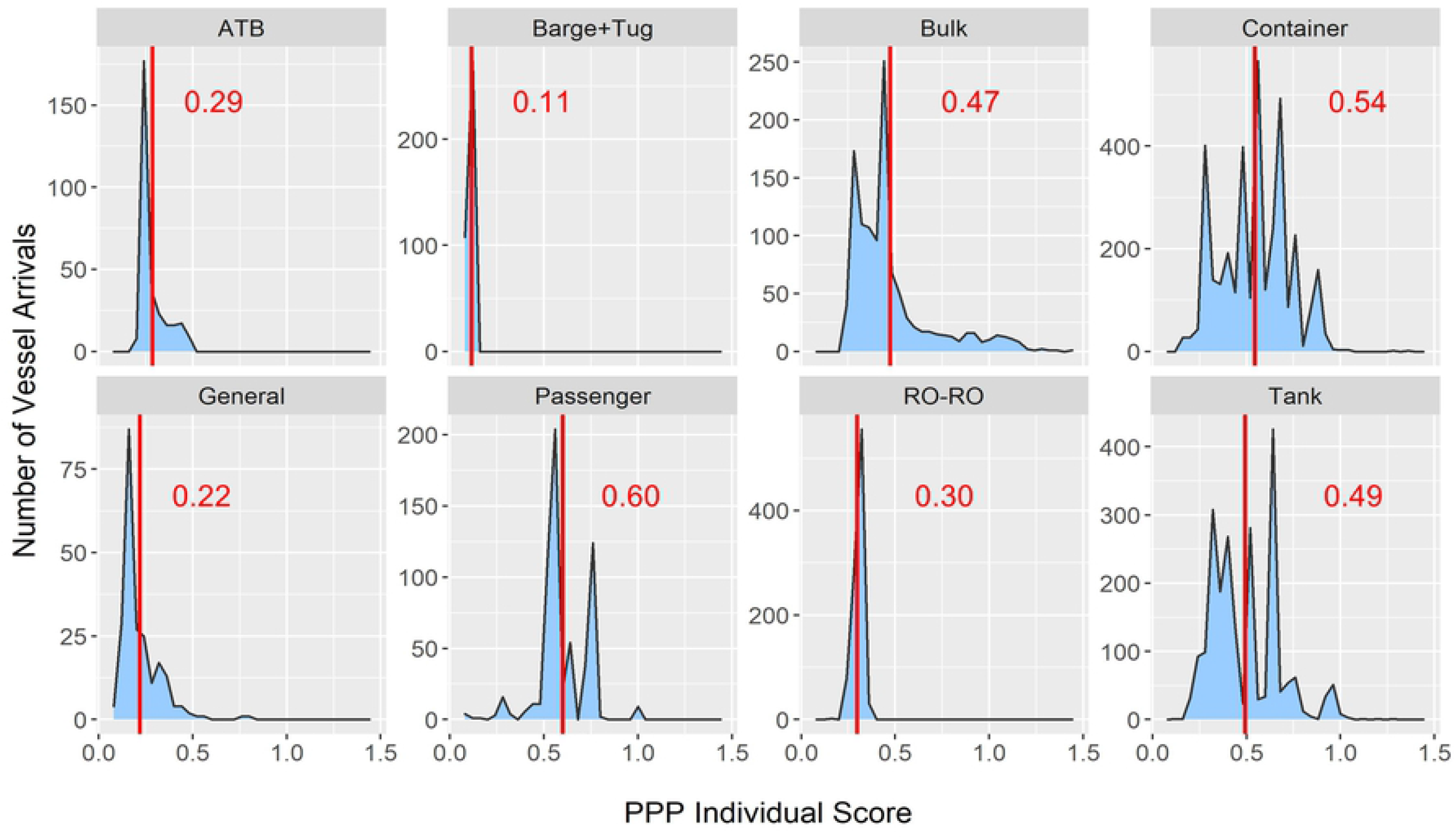
Frequency of PPP scores by vessel type. PPP score calculated for each vessel arriving at California ports during 2018 using the proposed proxy-based method to assess the combined effect of both biofouling and ballast water and the likelihood of species introductions associated to each arrival. Red line and numbers represent the mean PPP score for each vessel type. The PPP score ranges from 0 (lowest perceived likelihood of invasion) to 2 (highest perceived likelihood of invasion).

When evaluating how the PPP score could have been used to prioritize inspections on a daily basis over the entire set of 2018 arrivals (using the legislative mandate to inspect 25% of arrivals as a resources threshold to categorize high priorities), California would have prioritized container (57.2% of all high priorities), tank (20.9%), bulk (11.0%) and passenger (10.0%) vessels (Table 2). These vessel types presented the highest PPP scores daily, reflecting the large volumes of ballast water frequently discharged by these vessels in combination with their large TWSA values.

**Table 2.** Comparison of prioritization schemes. Vessels identified as high priority using the proposed PPP-based prioritization model (Projected) and actual vessels prioritized as high priority for inspection during 2018 using the existing California prioritizing scheme that relies on more resources and allows a more refined assessment as described in Fig 1.

|  | High priorities based on California's prioritization in 2018 |  | Projected high priorities (25% of total arrivals) based on PPP score |  |
| --- | --- | --- | --- | --- |
|  | Number of arrivals (unique vessels) | % of all high priorities | Number of arrivals (unique vessels) | % of all high priorities |
| ATB | 24 (6) | 1.8 | 0 | 0* |
| Barge+Tug | 39 (6) | 2.9 | 0 | 0* |
| Bulk | 472 (348) | 34.7 | 240 (123) | 11.0* |
| Container | 130 (106) | 9.6 | 1253 (271) | 57.2 |
| General | 85 (76) | 6.3 | 1 (1) | 0* |
| Passenger | 37 (20) | 2.7 | 238 (21) | 10.9 |
| RO-RO | 123 (92) | 9.0 | 0 | 0* |
| Tank | 450 (285) | 33.1 | 458 (152) | 20.9* |
| **Total** | **1360 (939)** |  | **2190 (568)** |  |

\*Indicates the vessel types that would have resulted in a decrease in their relative proportion of the overall high priorities when using the proposed model instead of the actual approach used.

When comparing the trial results of the proposed model with the actual California vessel inspection prioritization method used during 2018 (based on ballast water-related risk, outreach opportunities, and compliance history), the proposed prioritization scheme would have resulted in more container and passenger vessels categorized as high priority for inspection and fewer (including zero in some cases) bulk, tank, general, ATB, RO-RO, and unmanned barges (Table 2). These differences were expected because the approach used in California in 2018 relied on additional factors and resources, allowing a more reliable and refined assessment that resulted in a broader distribution of inspections across vessel types (Fig. 1)

In the trial, the proposed prioritization scheme assumes that the greatest 6 individual PPP scores per day will be prioritized as high priority for inspection, regardless of whether the same vessel makes repeated visits. Other jurisdictions that use this approach should decide how often to inspect vessels that frequently arrive and maintain a good compliance record. For example, we reexamined our dataset to only include the first high priority arrival for each vessel during 2018. All other arrivals from the same vessel were removed from prioritization to avoid repeatedly inspecting the same vessels. This resulted in a severely reduced number of unique container, tank, and passenger vessels categorized as a high priority (Table 2). Reducing the number of high priorities in these groups, would release additional resources that could be used to include other components in the prioritization scheme (e.g., outreach).

#### Cumulative PPP scores (by location or vessel type)

Cumulative PPP scores calculated during the trial showed that the Los Angeles/Long Beach port complex and the Port of Oakland received the greatest PPP (Fig 3), primarily because of the frequency of containers and tank vessels that arrive at these ports.

**Fig 3.**
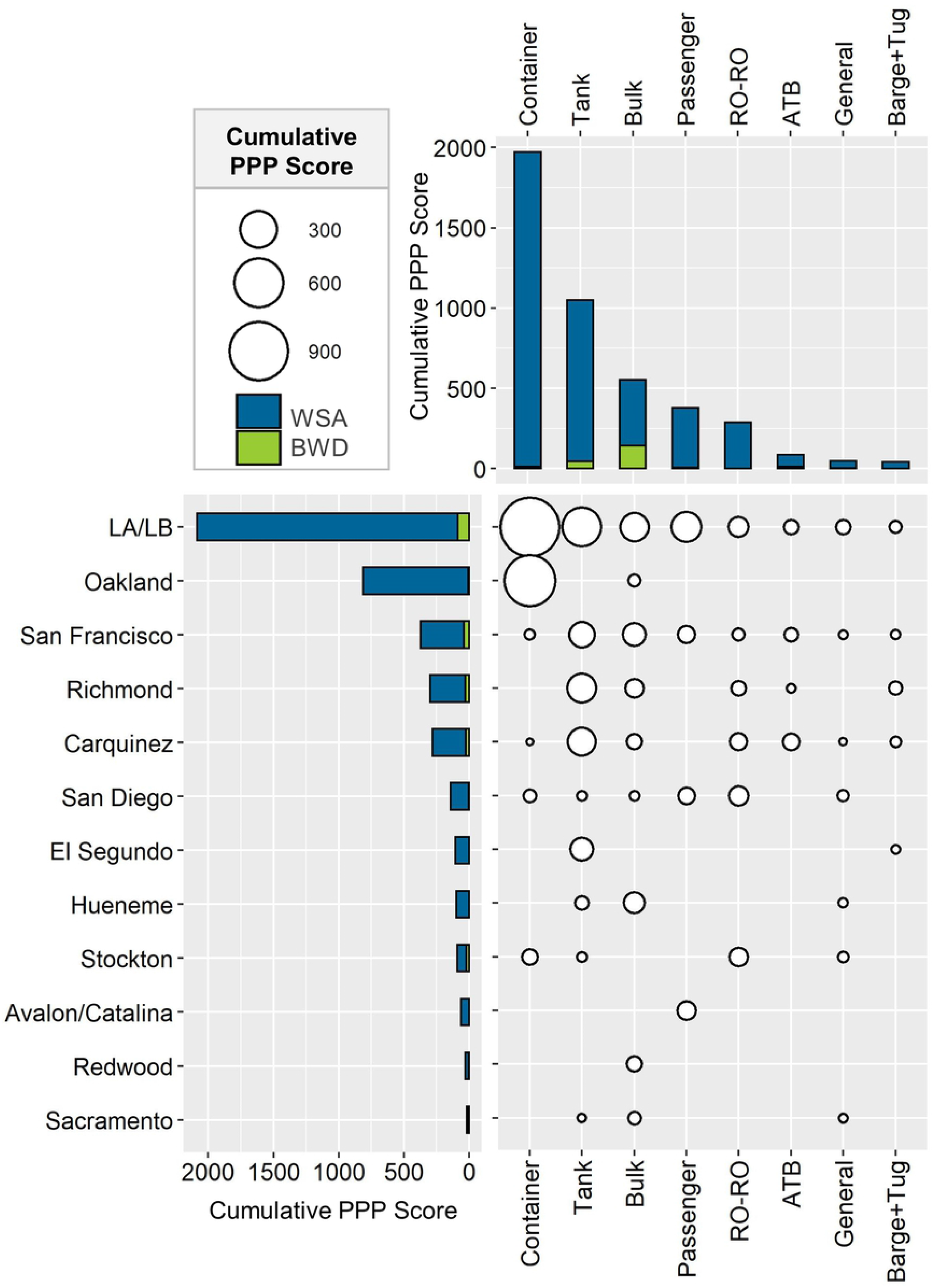
Cumulative PPP score observed in California during 2018. Bars represent the contribution of total wetted surface area (TWSA; blue) and ballast water discharge (BWD; green) to the combined cumulative score at all locations (horizontal) and all the different vessel types (vertical). Bubble size represents the cumulative score ranging from 1116.8 (largest bubble) to <1 (smallest bubble).

Across all ports and vessel types, TWSA outweighed BWD as a factor in the overall PPP scores, with BWD contributing to the scores primarily for tank and bulk vessels (Fig 3), reflecting the fact that only approximately 15% of all arrivals in California discharge ballast water [19].

## Discussion

The proxy-based model described here is a data-driven tool to prioritize ballast water and biofouling inspections of commercial merchant and passenger vessels. The approach is:

- Adaptable because it can be applied within any jurisdiction and relies on the specific characteristics of that jurisdiction’s vessel population
- Scalable because it can be adjusted to capture different segments of the vessel population (e.g., 10%, 15%, 25% of the arriving vessels)
- Adjustable because it can be altered to focus specifically on either ballast water or biofouling instead of taking a combined approach
- Versatile because scores can be added within groups (e.g., vessel types, different ports) or over time periods to differentiate the relative PPP within these groupings
- Simple because it can be set up in minutes and provide data-driven prioritization with only two readily available input values per arriving vessel

In the described form, this model is ideal for programs that regulate the management of both ballast water and biofouling as it considers the combined effect of proxies for both. However, it can be altered into separate or standalone prioritization approaches for programs that are primarily interested in only ballast water (e.g., some U.S. states) or biofouling (e.g., GloFouling Partnerships member countries). The approach also provides jurisdictions with a tool to determine the most efficient use of limited inspection resources on a daily basis or after a retrospective analysis of the patterns. Daily arrivals can be prioritized to ensure the inspection of vessels most likely to introduce NIS (assuming compliance with local management requirements) when additional data are unavailable. More efficient geographic allocation of inspection resources across multiple ports or regions can also be determined after analyzing the patterns of cumulative proxy-based PPP in these areas.

This proxy-based model relies on knowing each arriving vessel’s BWD volume and gross tonnage, both readily available to most jurisdictions. Most regulatory programs require submission of ballast water source, management, and discharge activity. In the U.S., vessels are required to submit the Ballast Water Management Report (BWMR) to the NBIC for every arrival at a U.S. port. Under the 2018 Vessel Incidental Discharge Act, the NBIC now makes all electronically submitted BWMRs (approx. 99.5% of submissions) available to state programs immediately upon receipt at NBIC, prior to arrival in most cases.

Vessel gross tonnage data are accessible at many free vessel information websites (e.g., vesselfinder.com) and can be used with the regression equations presented here, specific to each vessel type (see Table 1), to estimate WSA. The inclusion of niche area WSA in TWSA provides a more realistic proxy for biofouling PPP, as niche areas are known hotspots for biofouling accumulation [18, 20, 21].

Our trial of the proxy-based model with a dataset of California vessel arrivals demonstrates the utility of the method. To meet California’s legislative mandate to inspect at least 25% of arriving vessels, the trial using the proxy-based prioritization scheme focused inspections on container, tank, passenger, and bulk vessels. All of these vessels either frequently discharge ballast water in California and/or have large TWSA. When compared to the vessels actually categorized as a high priority in California during 2018 (based on outreach opportunities, compliance history, and BWD), the proposed model de-emphasized inspection of bulk and tank vessels and excluded ATBs, barges, auto carriers, and general cargo vessels from the high priorities. Even though compliance history was not included in this trial, it is important to emphasize that California’s prioritization process considers compliance with state requirements a critical part of the risk assessment process, as well as other factors like outreach.

The trial also verified the usefulness of using cumulative PPP scores for different ports to demonstrate how inspection resources can be allocated to those areas exposed to the greatest likelihood of introduction. This is especially useful for jurisdictions with ports separated geographically where the same set of inspectors cannot cover all ports.

The proxy-based model presented here is a baseline method that, in the absence of more detailed data, allows the detection of those vessels that may have the greatest likelihood of introducing NIS. The model assumes that management of both vectors has been performed according to the local requirements as the first layer of protection against NIS introductions. In addition, this approach does not consider opportunities for outreach as part of the prioritization process nor does it highlight the value of inspecting lower-scoring arrivals (e.g., for violation follow-up, or distribution of information to new vessels). These are important additional considerations that should be included in any prioritization scheme when possible. The flexibility of this method allows each jurisdiction to define outreach, compliance, or frequency rules (e.g., target new vessels, decrease inspection frequency in response to compliance history, suspicion of potential violations) that can be incorporated into the prioritization approach to reach a more comprehensive distribution of the inspection efforts. The approach also provides the opportunity for each jurisdiction to focus more attention (efforts) on ballast water or biofouling, depending on their priorities.

## Conclusion

The proxy-based prioritization model presented and trialed here is an adaptable, scalable, adjustable, versatile, and simple tool to rapidly identify a subset of vessels for ballast water and/or biofouling inspections. The approach can be adopted globally, and is especially useful for jurisdictions without access to, or authority to collect, risk profiling data or direct measurements for all incoming arrivals (Fig 1).

## Acknowledgments

Nicole Dobroski, Jonathan Thompson, Michelle Pelka, and Susie Kropman reviewed draft versions of the manuscript. Jackie Mackay, Dalia Keroles, Craig Fultz, Pearl Tan, Nadia Day, Michelle Pelka, Eric Frisbee, Paulette Boyd, Mary Fakih, Blanca Garcia, Michele Wiebold contributed with the data management of the California dataset. Kevin Reynolds contributed with valuable information and knowledge about vessel engineering for the WSA calculations.

## Supporting Information

**S1 Dataset**. California vessels arrivals from 2015 to 2018

**S2 Flowchart**. Step-by-step process to use the proposed approach.

**S3 Script**. R Script to calculate TWSA and PPP score using the proposed approach.

**S4 Data frame template**. Excel data frame template to input “Population Data” and “Arrivals data”.

**S5 Script**. R Script used to generate the figures for a visual analysis of the data.

